# Causes of death among hospitalized adults with dengue fever in Tainan, 2015: emphasis on cardiac events and bacterial infections

**DOI:** 10.1101/507137

**Authors:** Jen-Chieh Lee, Cong-Tat Cia, Nan-Yao Lee, Nai-Ying Ko, Po-Lin Chen, Wen-Chien Ko

## Abstract

**Introduction:** The 2015 dengue outbreak in southern Taiwan caused substantial mortality rates in the elderly. We analyzed here the causes of death among adults with dengue.

**Methods:** The retrospective study was conducted at a medical center in Tainan from the 1^st^ of August to 31th of December in the year 2015. The detection of the dengue NS1 antigen IgM or viral RNA in patients’ blood were used to diagnose dengue. Clinical courses and causes of death were retrieved from chart reviews by two intensivists.

**Results:** There were 4,488 cases of dengue diagnosed in the study hospital, and these cases had an in-hospital case fatality rate of 1.34% (60 cases). Of these, the mean age was 73 years and gender did not predict outcome. Twenty-eight (46.7%) cases died of severe dengue, and 29 (48.3%) deaths were possibly caused by dengue-related complications, which were mostly secondary infections (24 cases). Most of the families of fatal case (70%) signed do-not-resuscitate (DNR) orders prior to the patients’ death. When the dengue epidemic peaked, 13 cardiac arrest events, including out-of-hospital (5 events) and in-hospital (8) cardiac arrests at the emergency department, occurred within four weeks of the dengue epidemic. Notably, in half (7) of these cases, the patients did not search for medical aid prior to experiencing cardiac arrest. Of the 40 cases that had early death (occurring within one week after hospitalization), 60% died of severe dengue. In contrast, 50% of the 20 deaths that occurred later than one week after hospitalization were related to hospital-acquired infections, mainly pneumonia.

**Conclusion:** The elderly that experience dengue fever may die of severe dengue early or die of secondary infections later. Cardiac arrests can also occur unpredictably at the first aids, which highlight the need of professional and patient education regarding the danger signs that are related to severe dengue in an epidemic setting.

**Author summary:** The 2015 dengue outbreak in Tainan City caused substantial deaths among the elderly. The main causes of death were severe dengue and its complications. We here highlight the deaths caused by heart complications of dengue that the elderly has underlying cardiovascular comorbidities is more prone to be involved. The presentations of heart complications vary, ranging from arrythmia to myocarditis and to unexpected cardiac arrest. Clinicians should carefully evaluate and monitor the heart function of patients with severe dengue and provide timing intervention. Secondary infections or healthcare-associated infections may occur throughout the whole hospitalized course. They were also the leading causes of death during the late or recovery phase of dengue in the study. Judicious application of antimicrobial agents and early elimination of infection source may be beneficial. Overall, the substantial deaths during this outbreak may be related to low public awareness of dengue, emergency department overclouding, and lack of clinical experience. Professional and public education regarding the danger signs that are related to severe dengue is necessary in an epidemic setting.

## Introduction

Dengue is a mosquito-borne viral disease that affects humans and is emerging as a major threat to public health throughout the tropics and subtropics [1, 2]. Dengue affects children primarily in hyper-endemic areas [3] where epidemiological studies have shown the susceptible age group moved toward older children and adults [4]. Taiwan is not a hyperendemic area, but dengue outbreaks develop almost every year [5, 6]. Dengue predominantly affects adults in Taiwan. The highest prevalence rate is among people in their sixties [6]. During 2015, a large-scale dengue outbreak, which was caused by the dengue virus serotype 2, occurred in Tainan City [7] and a total of 22,777 dengue cases were confirmed [8]. Among these, 2%-3% had severe diseases or complications that required intensive care [9, 10]. However, with the increased utilization of intensive care units, the hospital stays for severe dengue patients were prolonged and more invasive devices were used. Healthcare-associated infections thus became unavoidable during the recovery phase.

Atypical presentations of dengue in the elderly are not uncommon [11] and make early diagnoses difficult. With multiple comorbidities complicating the clinical course of dengue, the elderly has more severe presentations and a higher mortality rate than children and young adults during this outbreak [12]. To know their cause of deaths may alarm early symptoms and signs of life-threatening conditions to clinicians. However, this information was limited in the literature. A two-year review in Malaysia has discussed deaths that were directly caused by dengue [13]. The causes of death stratified by the pathophysiology of dengue fever include dengue shock syndrome, severe bleeding, and severe organ involvement. Two case series that include seven and ten cases, respectively, mention deaths that were related to the complications other than dengue [14, 15]. Other studies that have paramedical points of view, discuss how these factors involve clinicians, patients, and the medical system, will lead to dengue death [16, 17]. The present study aimed to analyze the causes of death among dengue patients that were cared at a medical center in 2015.

## Methods

### Case inclusion

The study was conducted at the National Cheng Kung University Hospital (NCKUH), a medical center in southern Taiwan, between August 1^st^ and December 31^th^ in the year 2015. Cases of dengue were identified from the dengue notification in the Infection Control Center, NCKUH. The diagnosis of dengue fever was established by the documentation of dengue NS1 antigen (Bioline Dengue NS1 Ag kit, Standard Diagnostics Inc., Korea), dengue IgM (Bioline Dengue Duo kit, Standard Diagnostics Inc., Korea), or dengue virus RNA (TIB Molbiol, Lightmix kit, Roche Applied Science, Berlin, Germany) in the sera or blood of the patients. The study protocol was approved by the Institutional Review Board at NCKUH (A-ER-104-386) and informed consents were waived. The study was funded by Ministry of Health and Welfare, Taiwan, with award number: MOHW107-TDU-B-211-123003. The funders had no role in study design, data collection and analysis, decision to publish, or preparation of the manuscript.

### Definitions

Cardiac arrest events at the emergency department (ED) included out-hospital cardiac arrest (OHCA) and in-hospital cardiac arrest (IHCA). The weekly mortality rate was determined as the ratio of fatal case number and the number of newly diagnosed cases of dengue fever during the indicated week. Typical presentations of dengue fever included fever with two of the following symptoms or signs: nausea/vomiting, rash, aches/pains, positive tourniquet test, leucopenia, or any of warning signs [18]. If the initial manifestations of dengue did not fulfil the above criteria of the typical presentations, these cases were considered to be atypical. The severity of dengue fever was graded using the WHO guidelines issued in 2009 [18]. Group A includes those without warning signs, Group B those with warning signs, co-existing complicated conditions, or special social circumstances, and Group C includes those with the signs of severe plasma leakage, severe hemorrhages, or severe organ impairment. Plasma leakage was supported by a hematocrit change of more than 20% or the presence of pleural effusion or ascites. Unstable vital signs indicate the presence of hypotension (systolic pressure less than 90 mmHg or mean pressure less than 65 mmHg), impending respiratory failure, or altered mental status. The age-modified Charlson’s index was used to evaluate comorbidity status [19]. Days of hospitalization here were included days of ED stay.

Organ failure associated with dengue fever was defined as the dysfunction of the cardiovascular system, respiratory system, nervous system, liver, or kidney after dengue onset. Cardiovascular failure was defined as the patients having a mean blood pressure of less than 65 mmHg, the need of inotropic agents to maintain blood pressure, or low cardiac output signs. Respiratory failure was defined as a PaO_2_/FiO_2_ ratio of less than 300 [20] or PaCO_2_ more than 50 mmHg [21], with or without increased breathing effort. Nervous system failure was characterized as the patient having a Glasgow Coma scale score of less than 8 [22]. Serum bilirubin level of more than 2 mg/dl was regarded as liver failure [20]. The KDIGO criteria were applied to define renal failure, *i.e*., an increase of serum creatinine levels of more than twice the baseline level or more than 4 mg/dl, urine output of less than 0.3 ml/kg/hour over 24 hours, or anuria for 12 hours [23]. Multiorgan failure was defined as the presence of failure of more than two organ systems.

### Causes of death

Cause of death in the cases of dengue fever was evaluated by two intensivists with complete training in infectious diseases. The causes of death were categorized into three groups. Death was considered to be related to “severe dengue” if the patient died of severe plasma leakage, major bleeding, or organ failures. If the patient died of organ failures but we were unable to attribute this death solely to severe dengue, the death was classified as “possibly dengue related”. If there were other clinical diseases that inevitably lead to death, the cause of death was classified as “other”.

### Data acquisition

All parameters including demographic information, underlying disease, status of organ system dysfunction, date and cause of death, dengue onset, date of hospital visiting, and timing of the do-not-resuscitation (DNR) order, were obtained and calculated from electronic medical records.

### Statistical methods

To analyze the causes of death, patients were further stratified into deaths within one week of hospitalization and deaths after one week. Chi-squared or Fisher exact tests were used to compare categorical variables between groups. Continuous variables were compared using a Student’s t-test. All data were processed using SPSS (20th edition, IBM). A *P* value of less than 0.05 is denoted as significant.

## Result

During the study period, there were 4,488 hospitalized cases of serologically documented dengue in NCKUH, Tainan. Among these patients, 60 died with a crude in-hospital mortality rate of 1.34%, and 40 (66.7%) died within seven days of hospitalization. The mortality rate reached a peak of 1.57% during the 40th week and the case number reached its peak in the 36th week of 2015.

The mean age of patients in fatal cases was 73 years and the major underlying medical illnesses were hypertension (32 cases, 53.3%), diabetes mellitus (30, 50%), and chronic kidney disease (20, 33.3%), as shown in Table 1.

**Table 1.** Clinical characteristics of 60 fatal cases of dengue fever.

| Parameters | Case No. (%) |
| --- | --- |
| Age (mean ± standard deviation), years | 73.4 ± 12.4 |
| Underlying disease |  |
| Hypertension | 32 (53.3) |
| Diabetes mellitus | 30 (50.0) |
| Chronic kidney disease | 20 (33.3) |
| Cancer | 16 (26.7) |
| Coronary arterial disease | 15 (25.0) |
| Neurological disease | 12 (20.0) |
| Rheumatic disease | 8 (13.3) |
| Congestive heart failure | 7 (11.7) |
| Chronic hepatitis | 7 (11.7) |
| Liver cirrhosis | 4 (6.7) |
| Chronic obstructive pulmonary disease or asthma | 4 (6.7) |
| Chronic renal replacement therapy | 2 (3.3) |
| Recent chemotherapy | 2 (3.3) |

Neurological disease indicates one or more of old stroke, dementia, parkinsonism, or multiple sclerosis.

As for the causes of death, 46.7% (28 cases) died of severe dengue, and 65% (39) of the deaths were possibly related to dengue (Table 2). Of the later, a substantial proportion (40%, 24 cases) died of secondary infections. Two (3.3%) deaths were related to acute renal failure, and two (3.3%) died of respiratory failure without advanced airway management. A patient that refused interventional procedures died of massive gastrointestinal bleeding, which occurred while the patient had a platelet count of 94×10^9^/L during the 22^nd^ day of hospitalization. Three patients died from a brain tumor, medication suffocation, and a traffic accident that caused a skull bone fracture and intracranial hemorrhage and were coincident with dengue fever, respectively.

**Table 2.** Clinical characteristics of the deaths of adults with dengue fever occurring ≤1 week and >1 week after their hospitalization.

| Variables | Death after hospitalization |  | *P*<br>value |
| --- | --- | --- | --- |
| | $\leq 1$ week, n=40 | $>1$ week, n=20 | |
| Age (mean ± standard deviation), years | 73.1 ± 13.4 | 75.2 ± 10.5 | 0.52 |
| Male gender | 17 (42.5) | 13 (65) | 0.17 |
| Causes of death |  |  |  |
| Severe dengue | 24 (60) | 4 (20) | 0.03 |
| Intracranial hemorrhage | 2 (5) | 3 (15) | 0.32 |
| Cardiac events | 5 (12.5) | 0 (0) | 0.16 |
| Possibly dengue-related | 14 (35) | 15 (75) | 0.003 |
| Secondary infections | 14 (35) | 10 (50) | 0.58 |
| Bacteremia | 6 (15) | 0 (0) | 0.17 |
| Candidemia | 2 (5) | 1 (5) | 1 |
| Pneumonia | 5 (12.5) | 8 (40) | 0.02 |
| Pneumonia and bacteremia | 1 (2.5) | 0 (0) | 1 |
| Necrotizing fasciitis | 0 (0) | 1 (5) | 0.33 |
| Renal failure and its complication | 0 (0) | 2 (10) | 0.11 |
| Respiratory failure | 0 (0) | 2 (10) | 0.11 |
| Lower gastrointestinal bleeding | 0 (0) | 1 (5) | 0.33 |
| Others | 2 (5) | 1 (5) | 1 |
| Do-not-resuscitate | 24 (60) | 18 (90) | 0.02 |
| Cardiac arrest events at ED | 13 (32.5) | 0 (0) | 0.03 |

ED, emergency department

The death causes were compared between those dying within one week and those after one week of hospitalization (Table 2). For the former, the primary causes of death were severe dengue (24 cases, 60%) and secondary infection (14, 35%). Of the 24 deaths caused by severe dengue, five (12.5%) died from cardiac events that included fatal arrhythmia, myocarditis, and refractory heart failure. Among those that died after one week of hospitalization, the major causes of death were secondary infections (10 cases, 50%), renal failure (2, 10%), and respiratory failure (2, 10%). There were four (20%) cases in which the patients died from severe dengue. Of these, three (15%) died from intracranial hemorrhage, which developed during the critical phase of dengue. Candidemia can occur either early (within one week, two cases) or late (after one week, one case) during hospitalization. In contrast, among the seven fatal cases in which death was caused by bacteremia, all of these deaths occurred during the first week.

The mortality rate and the number of new cases and cardiac arrest events at the ED by week are shown in Figure 1. Of note, 13 cardiac arrest events occurred between the 36th and 39th week of 2015, also when the dengue outbreak peaked. Among the cases that had cardiac arrest, five occurred out-of-hospital cardiac arrest (OHCA) and eight had in-hospital cardiac arrest (IHCA) while staying at the ED (Table 3).

**Figure 1.**
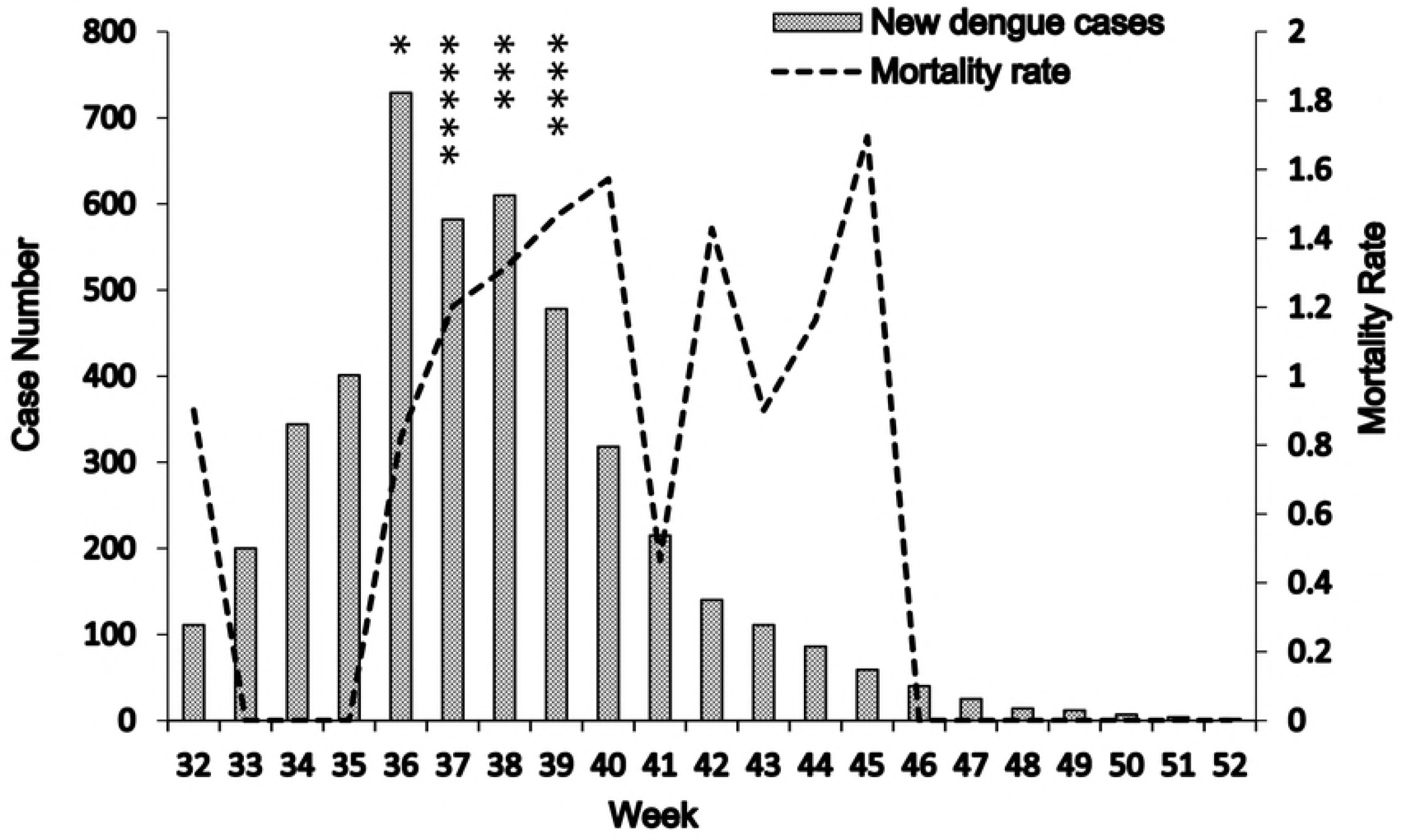
Weekly notified dengue cases in the study hospital and early mortality rates. The case number of newly diagnosed dengue patients came to peak in the Week 36 while the early mortality rate (*i.e*., death within the first week of hospitalization). peaked in the Week 40. Cardiac arrest events occurred mainly in Week 36-39 when the outbreak was in full swing. An asterisk (*) indicates a cardiac arrest event at the emergency department.

**Table 3.**
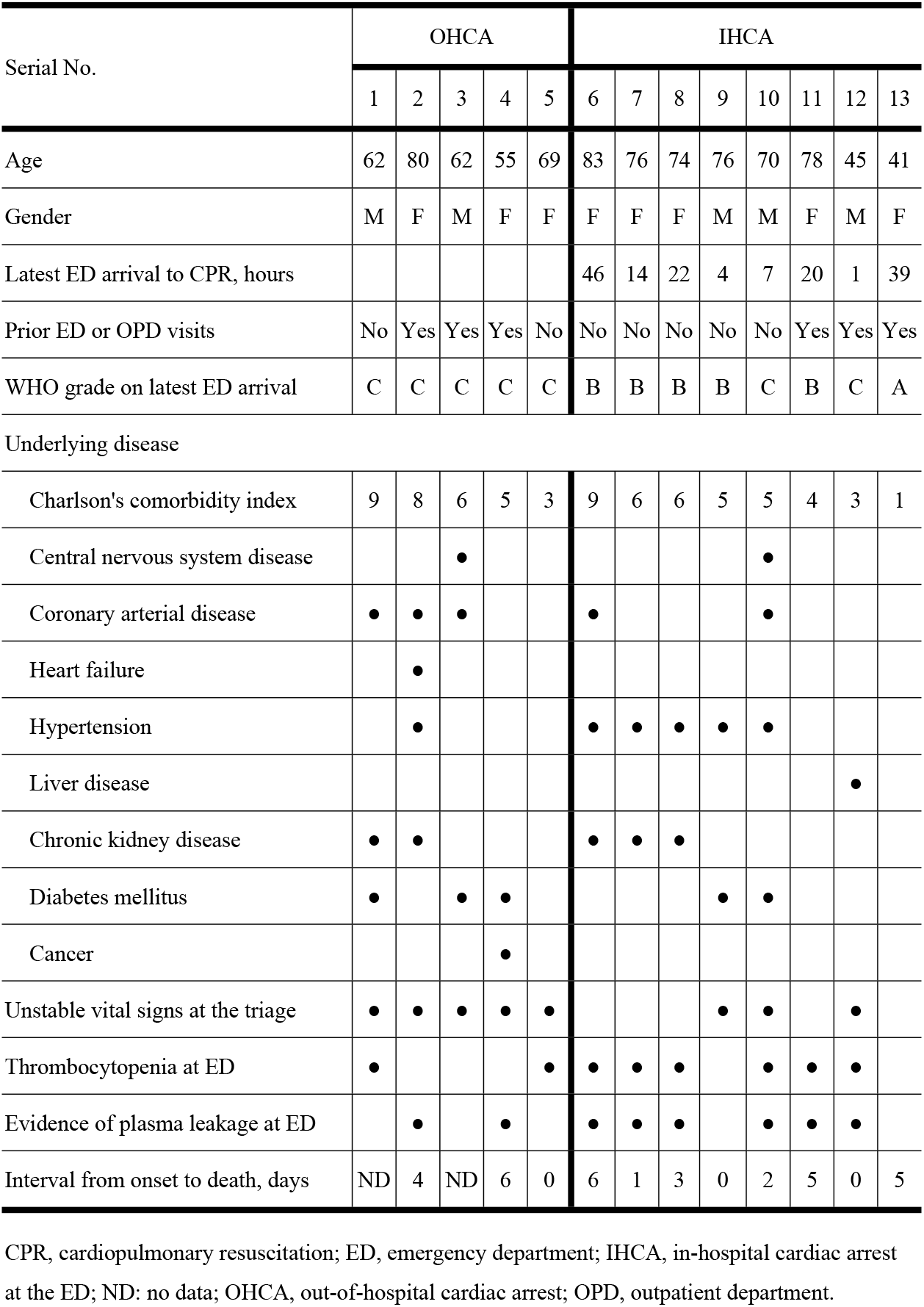
Characteristics of 13 cardiac arrest events at the emergency department.

| Serial No. | OHCA |  |  |  |  | IHCA |  |  |  |  |  |  |  |
| --- | --- | --- | --- | --- | --- | --- | --- | --- | --- | --- | --- | --- | --- |
|  | 1 | 2 | 3 | 4 | 5 | 6 | 7 | 8 | 9 | 10 | 11 | 12 | 13 |
| Age | 62 | 80 | 62 | 55 | 69 | 83 | 76 | 74 | 76 | 70 | 78 | 45 | 41 |
| Gender | M | F | M | F | F | F | F | F | M | M | F | M | F |
| Latest ED arrival to CPR, hours |  |  |  |  |  | 46 | 14 | 22 | 4 | 7 | 20 | 1 | 39 |
| Prior ED or OPD visits | No | Yes | Yes | Yes | No | No | No | No | No | No | Yes | Yes | Yes |
| WHO grade on latest ED arrival | C | C | C | C | C | B | B | B | B | C | B | C | A |
| Underlying disease |  |  |  |  |  |  |  |  |  |  |  |  |  |
| Charlson's comorbidity index | 9 | 8 | 6 | 5 | 3 | 9 | 6 | 6 | 5 | 5 | 4 | 3 | 1 |
| Central nervous system disease |  |  | • |  |  |  |  |  |  | • |  |  |  |
| Coronary arterial disease | • | • | • |  |  | • |  |  |  | • |  |  |  |
| Heart failure |  | • |  |  |  |  |  |  |  |  |  |  |  |
| Hypertension |  | • |  |  |  | • | • | • | • | • |  |  |  |
| Liver disease |  |  |  |  |  |  |  |  |  |  |  | • |  |
| Chronic kidney disease | • | • |  |  |  | • | • | • |  |  |  |  |  |
| Diabetes mellitus | • |  | • | • |  |  |  |  | • | • |  |  |  |
| Cancer |  |  |  | • |  |  |  |  |  |  |  |  |  |
| Unstable vital signs at the triage | • | • | • | • | • |  |  |  | • | • |  | • |  |
| Thrombocytopenia at ED | • |  |  |  | • | • | • | • |  | • | • | • |  |
| Evidence of plasma leakage at ED |  | • |  | • |  | • | • | • |  | • | • | • |  |
| Interval from onset to death, days | ND | 4 | ND | 6 | 0 | 6 | 1 | 3 | 0 | 2 | 5 | 0 | 5 |

CPR, cardiopulmonary resuscitation; ED, emergency department; IHCA, in-hospital cardiac arrest at the ED; ND: no data; OHCA, out-of-hospital cardiac arrest; OPD, outpatient department.

The intervals between ED arrival and the IHCA event varied, ranging between one and 46 hours. The estimated interval between dengue onset and death ranged between 0 and 6 days. Ages in these cases ranged between 41 and 83 years and the Charlson’s index ranged between 1 and 9. Of these 13 cases, 7 cases did not search for medical aid prior to the events, and 12 cases presented to the ED with severe dengue (WHO Grade B or C). The exception was a case of WHO Grade A disease that died soon after the onset of upper back pain with diffuse ST-T elevation (Figure 2) in electrocardiography records at the ED. This is shown as case 13 in Table 3.

**Figure 2.**
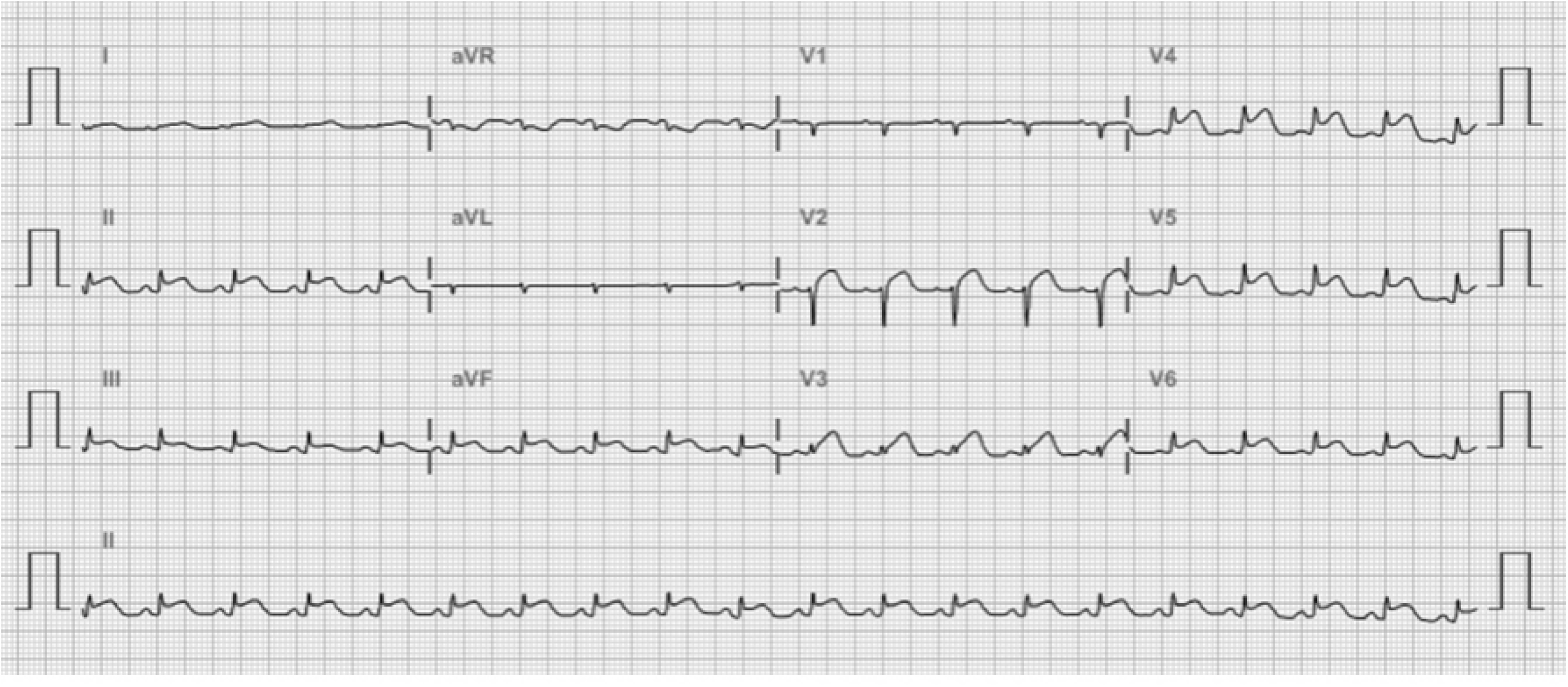
Electrocardiography record from a patient with dengue fever that was dying of myocarditis. The diagnosis of myocarditis was suggested by the diffuse ST-T elevation (case 13 in Table 3).

Of the 60 fatal cases, 70% (42) signed do-not-resuscitate (DNR) orders prior to their death (Table 4), and two thirds (28 cases) of DNR decisions were made within one week after their hospital arrival. Of note, DNR was requested in six cases without organ failure, twelve with failure of one organ system, and eight with failure of two organ systems

**Table 4.** Number of organ failures in the 42 fatal cases of dengue fever at the time of the do-not-resuscitate (DNR) signature.

| Parameters | Case No. (%) |
| --- | --- |
| No organ failure | 6 (14.3) |
| Failure of one organ system | 12 (28.6) |
| Respiratory failure | 5 (11.9) |
| Renal failure requiring renal replacement therapy | 4 (9.5) |
| Cardiovascular failure | 2 (4.8) |
| Central nervous system (CNS) failure | 1 (2.4) |
| Failure of two organ systems | 8 (19.0) |
| Renal and respiratory failure | 3 (7.1) |
| CNS and respiratory failure | 3 (7.1) |
| Cardiovascular and respiratory failure | 1 (2.4) |
| CNS and renal failure | 1 (2.4) |
| Multiorgan failure | 16 (38.1) |

## Discussion

The 2015 dengue outbreak in southern Taiwan came to its peak in early September. The mortality rate of patients included in this study reached its peak during Week 40. Of note, the majority of the sudden death events, including OHCA and IHCA at the ED, occurred within the first four weeks of the dengue outbreak. This is likely related to the following: first, public awareness of dengue was low during the early phase of the epidemic and less than one half (6) of the 13 patients that experienced cardiac arrest sought medical care before their death. Early medical care could be life-saving before dengue deteriorated [13-15]. Second, ED overclouding may be associated with increased mortality rates and morbidity [24, 25], and the role of patient diversion [24, 26] and the short stay ward at the ED [27] during large outbreaks of infectious diseases is mentioned. With the implementation of a patient diversion program in Tainan, and dengue rapid screening clinics in the study hospital, the ED patient burden was correspondingly decreased, which may facilitate optimal initial care for the elderly that have severe dengue. Third, a lack of clinical experience in the ED professional staff may delay the timely diagnosis and appropriate care for severe diseases. A recent study found that case management training can improve dengue outcomes [28] and protocolized care may accelerate the transformation of knowledge [29] and help inexperienced healthcare providers.

Dengue in Taiwan often involves the elderly [6], for which the diagnosis of dengue may be challenging because of atypical presentations [5, 30]. The febrile elderly with leucopenia has been referred to as a trigger for diagnostic laboratory tests for dengue fever [31]. The clinical symptoms and signs that are presumably linked to underlying comorbidities in the elderly can further delay the diagnosis of dengue. More than half of our patients had atypical presentation during their initial medical visits, and the diagnosis of infection with the dengue virus was delayed in eight cases. Thus, to achieve the goal of early diagnosis, clinical suspicion of dengue should not be limited to the typical dengue fever “criteria” in endemic settings.

In our series, five OHCA and eight IHCA cases that had atypical presentation at the ED had a fulminant course. Not all fatal dengue cases received optimal intensive care in the ED before their death. Thrombocytopenia was not observed in five cases, and five had no evidence of plasma leakage, contradictory to severe manifestations of dengue fever [18]. Closely laboratory test and vital sign monitoring may be necessary in patients really needed intensive care. Another potentially fatal complication of dengue fever is pericardiomyocarditis, which is illustrated by sudden collapse, diffuse ST-T elevation in electrocardiograms (Figure 2), and elevated serum cardiac enzymes, but without thrombocytopenia or plasma leakage.

The overall mortality rate of hospitalized patients with dengue fever in the study hospital was 1.3%. Multiple factors, such as increased age or epidemic serotype 2 dengue virus, may contribute to the substantial mortality rate in our study. One potential contributing sociocultural factor is the DNR signature during hospitalization in up to 70% of fatal cases. Through advances in organ support, those with severe dengue may have a greater chance to survive, but DNR orders precludes the intensive care provided by healthcare workers. In Taiwan, old age is the main factor in signing DNR orders in critically ill patients [32, 33]. A study in Taiwan revealed that patients with renal failure had the highest percentage of signed DNR orders [34]. Furthermore, aspects of Taiwanese culture caused many to refuse tracheostomy [35]. We observed that dengue patients with DNR orders and without organ failure were older than those with organ failures, although the difference was not significant. However, their prognosis may remain poor even without a DNR order under advanced medical support.

To further analyze the causes of death, more than half of the early deaths (*i.e*., deaths within one week of hospitalization) were related to severe dengue with either plasma leakage or major bleeding. However, there were three cases in which the patients experienced rapid organ failure without evidence of plasma leakage and major bleeding. In contrast, the two main causes of death that occurred after one week of hospitalization were healthcare-associated bacterial infections and organ dysfunction that was related to severe dengue. Identifying secondary infections among hospitalized dengue patients is critical during dengue epidemics. The primary pathogens that caused bacteremia in dengue patients were *Staphylococcus aureus, Streptococcus* species, and *Enterobacteriaceae* [36]. Old age, comorbidities, severe disease with gastrointestinal bleeding, prolonged activated prothrombin time, and acute renal failure were recognized as risk factors for concurrent bacteremia [37-39]. Laboratory parameters, such as leukocyte count [38], serum C-reactive protein [39], and procalcitonin [40], may be useful to differentiate those with concurrent bloodstream infections from those that had only dengue virus infection alone.

One unique finding in the present study is that cardiac events were associated with dengue virus infection. Before their death, five patients suffered cardiac events, including heart failure with cardiogenic shock or pulmonary edema, fatal arrhythmia, and fulminant myocarditis. In the literature, the disease spectrum for cardiac involvement in dengue patients was wide and ranged from subclinical functional myocardial impairment to fulminant myocarditis [41, 42]. Fluid resuscitation may predispose patients with impaired cardiac function to pulmonary edema, and arrhythmias may be induced when plasma leakage or renal function impairment was ongoing. Patients with risk factors, such as old age, underling cardiovascular disease, abnormal electrocardiogram findings, elevated cardiac enzymes, and shock unresponsive to fluid resuscitation, warrant further evaluation by cardiac specialists [43].

Our study has several limitations. First, this study was conducted at a single medical center in southern Taiwan, which provided care for more than half of the critical patients from Tainan City during the 2015 dengue epidemic. Second, the data were collected retrospectively, and certain important parameters, such as the time of disease onset and detailed symptoms and signs, cannot be obtained from medical records. Third, the causes of death were clinically categorized and cannot be verified by autopsy. However, the study results were recorded by two experienced intensivists who had the Taiwan board of infection disease specialization.

In conclusion, elderly patients with severe dengue are at risk of fatal cardiac events and secondary infections. Absence of medical care among one half of adults with cardiac arrest reveals the importance of health education for the public during dengue epidemics.

## Acknowledgement

We thank Chih-Cheng Hsieh and Wei-Chieh Lin for their help in conceptualization of the study and preparing the manuscript.

## Supporting information

**S1 File. The data base of the 60 fatal cases in 2015 dengue outbreak in Tainan. S1 Checklist. STROBE Checklist**

